# Hierarchical development of dominance through the winner-loser effect and socio-spatial structure

**DOI:** 10.1101/2020.12.01.406017

**Authors:** E.S. van Haeringen, C.K. Hemelrijk

## Abstract

In many groups of animals the dominance hierarchy is linear. What mechanisms underlie this linearity of the dominance hierarchy is under debate. Linearity is often attributed to cognitively sophisticated processes, such as transitive inference and eavesdropping. An alternative explanation is that it develops via the winner-loser effect. This effect implies that after a fight has been decided the winner is more likely to win again, and the loser is more likely to lose again. Although it has been shown that dominance hierarchies may develop via the winner-loser effect, the degree of linearity of such hierarchies is unknown.

The aim of the present study is to investigate whether a similar degree of linearity, like in real animals, may emerge as a consequence of the winner-loser effect and the socio-spatial structure of group members. For this purpose, we use the model DomWorld, in which agents group and compete and the outcome of conflicts are self-reinforcing. Here dominance hierarchies are shown to emerge. In the model, we apply analytical methods previously used in a study on dominance in real hens including an analysis of behaviourial dynamics and network triad motifs.

We show that when in the complete model one parameter, representing the intensity of aggression, was set high, the model reproduced the high linearity and many patterns of hierarchical development typical of groups of hens. Yet, when omitting from the model the winner-loser effect or spatial location of individuals, this resemblance decreased markedly.

Our results demonstrate that the spatial structure and the winner-loser effect provide a plausible alternative for hierarchical linearity to processes that are cognitively more sophisticated. Further research should determine whether the winner-loser effect and spatial structure of group members also explains the characteristics of hierarchical development in other species.

## Introduction

Dominance hierarchies are a near universal pattern of social order in group-living animals. High dominance rank is supposed to be adaptive for access to resources and protection from predators (1–5). The hierarchy is often (near) linear in small groups of up to 10 individuals in a wide range of species, including mammals, fish, birds, crustacean and insects (6, 7). Yet, what proximate mechanism causes linearity is under longstanding scientific debate.

For the formation of a dominance hierarchy mainly two mechanisms have been proposed. First, prior attributes of individuals such as body size or personality characteristics, have been suggested to directly determine dominance rank. This theory is well supported by empirical data on pair-wise dominance interactions (8), but it has been rejected in a theoretical study because hierarchies are (near) linear and this would require difficult mathematical conditions, especially in larger groups (9–11). Second, the self-reinforcing outcomes of a fight may result in hierarchy formation (12). The self-reinforcing effect implies that, the winner of a dominance interaction becomes more likely to win again, whereas the loser becomes more likely to lose again, the so-called winner-loser effect (13–15).

The winner-loser effect is effective in a wide variety of species. Most evidence comes from contests in experimental studies of isolated pairs, rather than in groups (11). An exception is a study by Lindquist and Chase (16) of small groups of hens. Here the authors tracked the development of the dominance hierarchy with novel analytical methods. They showed that the hierarchy became highly linear and stable and it developed fast. Attacks in pairs of hens often occurred in series of attacks in the same direction (bursts) and those network states occurred most often that either contained an individual that dominated all others, or that comprised only triads that were transitive (shown in a triad motif analysis of the network).

Additionally, the authors mathematically represented three models of hierarchy development based on the winner-loser effect. Namely, the Bonabeau model (17), the Dugatkin model (15) and the Hemelrijk model, called DomWorld (18). However, Lindquist and Chase ignore that DomWorld is an individual-based model with a spatial representation of individuals (16). They investigated whether their mathematical abstractions reproduce some aspects of the hierarchy formation in hens. Since they did not sufficiently reproduced observations, they concluded that hens are likely aware of the group hierarchy and actively strive for it to become linear.

Their detailed description of the formation of dominance hierarchies in hens offers an opportunity to examine for the first time whether the DomWorld model in its complete form (including the spatial representation of groups of individuals) suffices to generate hierarchical patterns similar to those in hens despite the model’s cognitively simple rules (agents are not striving for linearity of the dominance hierarchy). We here investigate the importance of the spatial representation of interactions for the formation of a highly linear and stable hierarchy, because the spatial component of the model in combination with the winner-loser effect has formerly been shown to contribute to the generation of a wide variety of complex patterns of social interaction resembling those in primates, including many aspects of egalitarian and despotic dominance styles of various species of macaques (19, 20).

Therefore, the aim of the present study is to investigate how the winner-loser effect and the socio-spatial structure affect the development of the dominance hierarchy in groups in the model, DomWorld when we use the same analytical methods as Lindquist and Chase (16) did for their studies on hens.

## Methods

### The model

The computer model DomWorld (18) is an individual-based model in which agents move in infinite space. The agents have a tendency to group when other agents are far away and engage in dominance interactions when other agents are within their ‘personal space’. DomWorld is event-driven and does not have a representation of time. Dominance interactions between agents can be either won or lost. The outcome of a fight is self-reinforcing, such that the winner becomes more likely to win subsequent fights and the loser more likely to lose these. Throughout this article we will use the terms ‘*win’* and ‘*lose’* for the outcome of dominance interactions and ‘*initiation’* for starting a dominance interaction, also referred to as a fight.

A dominance interaction is mediated by dominance values (DOM) that represent each agent’s fighting power. The chance *W*_*i*_ of agent *i* to win a fight against agent *j* is determined by comparing its ratio of the *DOM* values to a number drawn from a random distribution (Eq. 1).

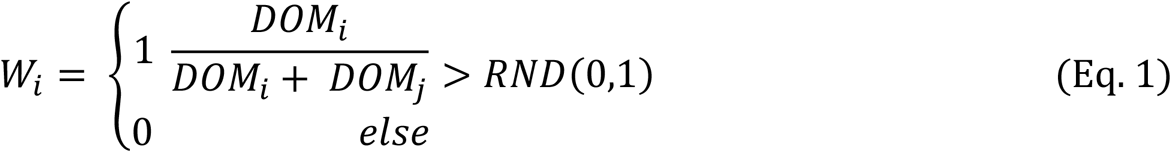

Afterwards the *DOM* values are updated depending on the outcome of the fight. The value of the winner (*DOM*_*i*_) increases with 1 minus its relative dominance ratio. The loser decreases its score by the same amount. The change in DOM value is multiplied by a scaling parameter *StepDom* that symbolises the intensity of aggression (Eq. 2).

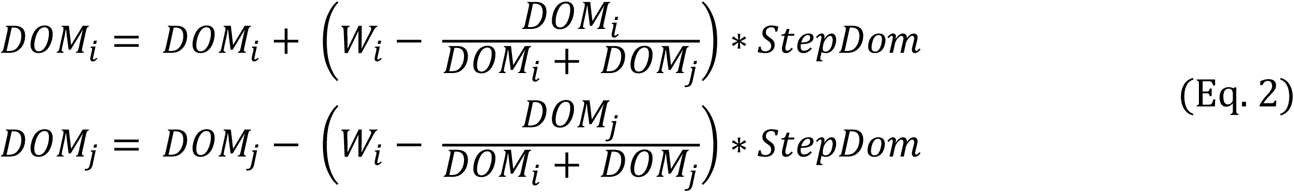

In the model, DomWorld, agents have been using two strategies of attack (18), obligate and risk-sensitive. In the present paper we will only discuss the obligate style of attack, as risk-sensitivity in general did not influence the patterns of hierarchy formation and was used in the work by Lindquist and Chase (16). When agents are meeting an individual in their personal space and are set to always attack it, this is referred to as an obligate strategy of attack (19). If an agent attacks only if it assumes it will win from the opponent, this has been called risk-sensitive attack. Here, an agent assumes it will win from the opponent if wins all mental fights, which it performs against a potential opponent, according to equation 1. A more extensive description of DomWorld can be found in Hemelrijk (21).

### Setup

Starting from the parameter setting from the work by Hemelrijk in 1999 (21), we tuned the model DomWorld to match several aspects of competition among real hens observed by Lindquist and Chase (16). We used the same number of groups (14), the same group size (4 females) and the same average number of interactions (518) as reported for the study of hens(8).

To obtain the same average linearity of the hierarchy in the model as reported in hens, we increased the value of a single parameter, *StepDom,* representing a higher intensity of aggression, thus, increasing the average linearity of the hierarchy in the model (Fig 1).

**Fig 1.**
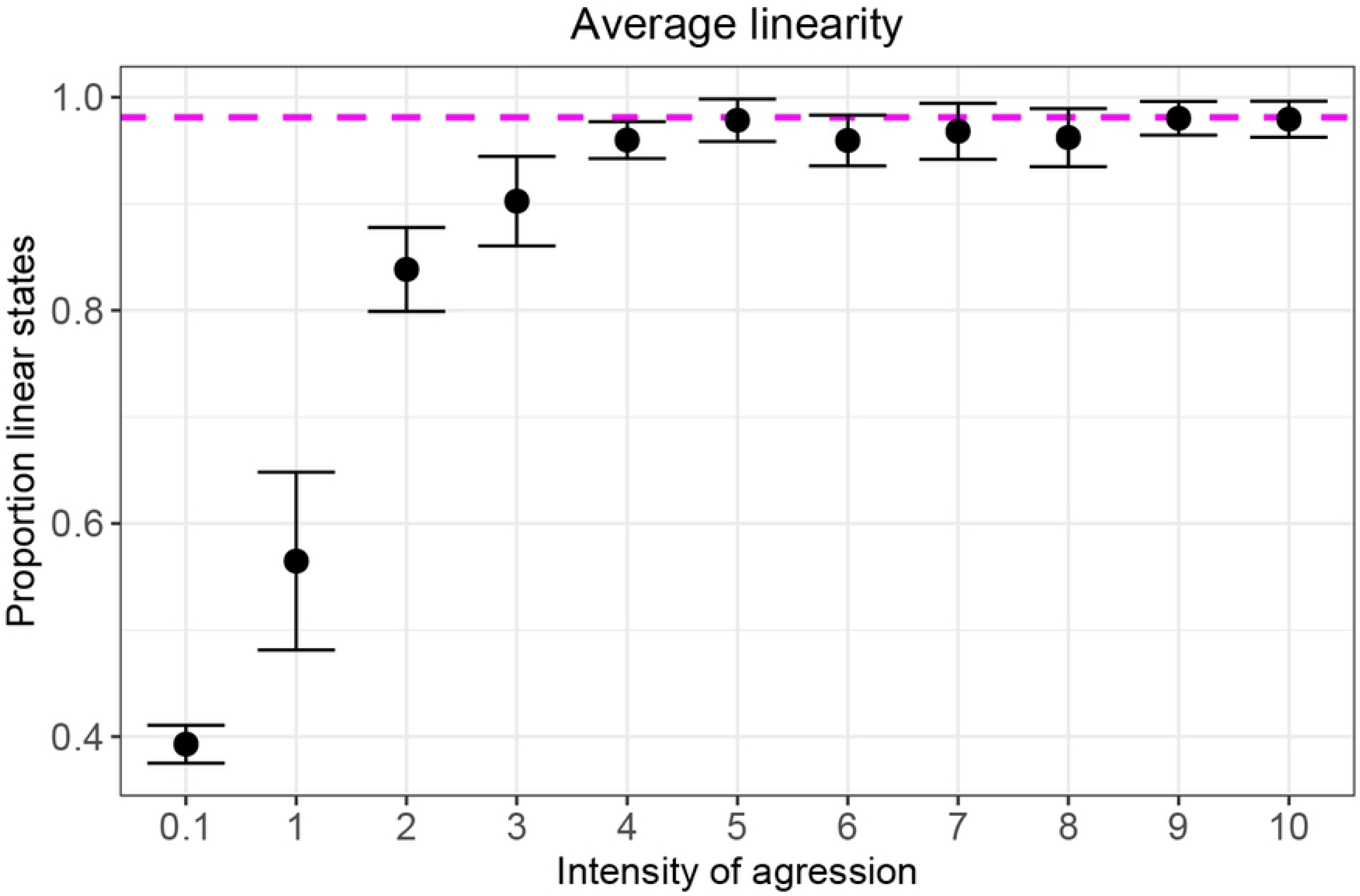
Average (and SE) of the hierarchical linearity measured as the proportion of transitive states for different levels of intensity of aggression. The dashed line indicates the proportion of transitive states estimated for the groups of hens in Lindquist and Chase (16).

We estimated linearity in the empirical data of hens from the study of Lindquist and Chase from the frequencies reported in their Fig 12 (16). Our estimation concerns the proportion of network states that is fully linear within the states with the same number of links. In 95% of the interactions in DomWorld the network contained the maximum of 6 links. Because in states with 6 links, state 38 is the only fully transitive state, we chose its proportion as our target value. It occurred in hens in 98% of the interactions with 6 links (see dashed line Fig 1). For the present paper we chose the lowest value of StepDom that matched this target, which is a value of StepDom of 9.

To gain understanding of the effects of spatial structure and the winner-loser effect we studied results not only in DomWorld in its complete form, but also in DomWorld without spatial structure (by selecting interaction partners at random) and without the winner-loser effect (by using fixed DOM scores of agents based on the final DOM scores of the simulations with the full model).

### Data collection

We calculated the various measures in our data-analysis using scripts written in Python (version 3.6.8), see the section *Analysis* below. The python package DHDAT of many of these measures is freely available, see (22). The statistical analysis and creation of figures were performed with R (version 3.5.1) in combination with packages Ggplot2, Multcomp and Dplyr.

### Analysis

In analysing the development of the dominance network in DomWorld we use methods and naming conventions similar to those by Lindquist and Chase in groups of hens (16).

### Patterns of dominance

We determine the hierarchy using the Average Dominance Index (ADI) which is the average of an individual’s proportion of wins from all its interactions against each interaction partner in the group (23). We calculated this cumulatively after each interaction. The differentiation of the hierarchy is measured with the steepness measure described by de Vries et al. (24) applied to the ADI of the agents in DomWorld. Note that the ADI is (almost) the same as the David score (23). We estimated the steepness of the hierarchy in the model with the same measure applied to the average dominance values reported by Lindquist and Chase (16) in figure 10. We used imaging software to sample 11 points per individual from this figure. Ten of these were taken from interaction 50 to 500 (one per 50 interactions). The first point was sampled after 10 interactions instead of 0, when there is no hierarchy.

Note that the hierarchy, and consequently all analysed metrics, is established using the ADI, based on the outcome of fights, not with the DOM scores of the agents in the DomWorld model, that symbolise their fighting power. Therefore, the order of the hierarchy can change in the model version without the winner-loser effect even though the fighting power (DOM) of the agents is fixed in this version.

We examine bursts and pair-flips. Bursts are defined as a series of at least two consecutive attacks in the same direction in a single pair of individuals. Important to note is that this definition does not involve a time interval. Attacks that are part of a burst can theoretically be widely separated in time, as long as there are no attacks involving other pairs in between. An attack in a dyad is labelled as a pair-flip when the direction of a subsequent attack is opposite to that of the previous one.

The development of the ADI per individual in the model we show as a separate horizontal line in a music notation graph (Fig 4A). A vertical arrow from the winner to the loser represents a directed dominance interaction. An ‘X’ at the top of the graph indicates a Pair-flip.

The degree of spatial centrality of dominant individuals in the model we measured for each activation by correlating the dominance rank of each individual with its distance to the centre of the group.

### Patterns in network structure

The development of the dominance network we describe with triad motifs (25, 26). Each motif comprises 3 individuals (nodes) and their 3 dyadic relationships (links or edges). Each triad in the network is labelled according to the presence and direction of dominance relationships among its members. When considering only directed relations, there are 7 triad motifs (Fig 2). If each individual has a directional relationship with two members of the triad, the triad can either be transitive (Fig 2F) or intransitive (Fig 2G). Transitive implies that A dominates both B and C, B dominates C, and C is subordinate to both. Cyclic or intransitive implies that while A dominates B, B dominates C and C dominates A.

**Fig 2.**
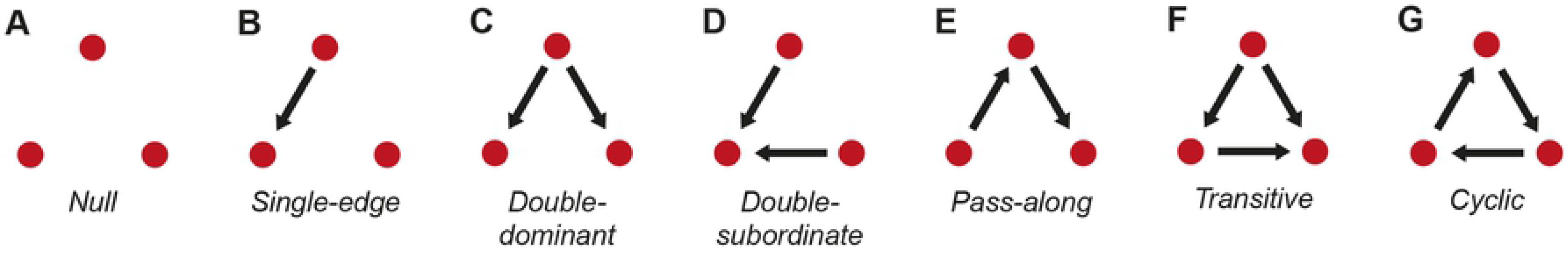
Motifs of directed relations in a triad. Motifs A to E are partial triads, motifs F and G are complete triads. Individuals in motif F can be ranked. Therefore, motif F is transitive. Individuals in motif G cannot be ranked and therefore, it is intransitive (cyclic).

In a directed network with 4 individuals there are 4 triads and 41 different states of these 4 triads, plus 1 state with no relations among the individuals, which is not included (Fig 3). States are categorised in groups with the same number of dyadic relations in the network, the so-called link-group. In a link-group states are categorised in classes (indicated by a letter). States in the same class share a network structure such that a pair-flip can change the network state to another one in the same class, but not to a network state in a different class (or link-group). If there is an individual that is dominant over all others (indicated as DAO) in the group, its node is marked with a ‘D’. The node of an individual that is submissive to all others (SAO) is marked with an ‘S’. The development of the network is traced over time by recording its state after each interaction.

**Fig 3.**
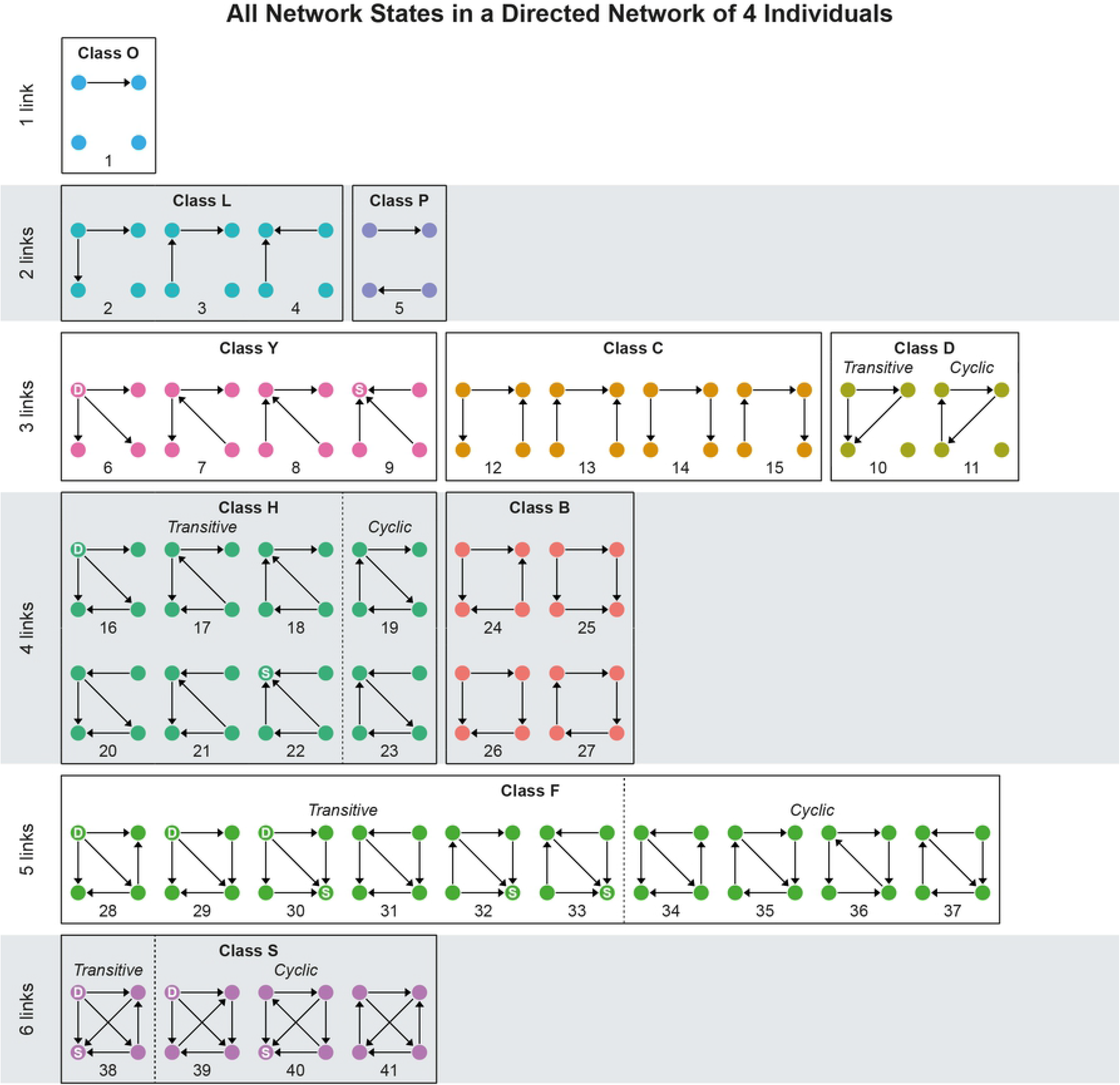
All states of a directed network of 4 individuals. The states are categorised by number of links (edges), by class and transitivity, following (16). Note that the colours of the classes correspond to the colours in Fig 7. A node marked with a ‘D’ is an individual that Dominates All Others (DAO), and marked with an ‘S’ is an individual Submissive to All Others (SAO).

The degree of transitivity is measured as the proportion of states that are completely transitive (without any cyclic triad) out of all states with at least one complete triad. We also calculated the transitivity, T_tri_, as described by Shizuka and McDonald (26). It is the proportion of complete triads that are transitive, normalised by the proportion transitive triads that are expected on average in a random network. Because a state can also be partially transitive, in theory this proportional measure of transitivity has a higher resolution than its binary definition of a state being either completely transitive or cyclic. Since results for the binary and proportional definition of transitivity were similar for the settings in the present paper, we only show the proportion of transitive states, T_tri_.

The occurrence of each network state is examined using two measures: the Class Occurrence Frequency (COF), which is the proportion of simulations in which a state occurred at least once, and the Class Stability Frequency (CSF), which is the number of interactions that occurred while the dominance network was in a particular state, divided by the total number of interactions that occurred in all the states with the same number of links. For each state, the average Class Occurrence Frequency, COF, and the Class Stability Frequency, CSF, are shown in a histogram categorised according to the link-group and the class with the number of recorded interactions to indicate the degree of accuracy (Fig 7).

### Statistical analysis

We tested the effects of removing space from the model and of removing the winner-loser effect using a generalized linear model (GLM) for each measure in Table 1. Other predictor variables included the identifier of the run, and for rank-changes that correlated with pair-flips (item 2 in Table 1) rank-changes and pair-flips were included as predictors. For analysing hierarchy steepness, the maximum length of bursts and the proportion pair-flips and rank-changes during early hierarchy formation (items 3, 4, 6 and 8 of Table 1) we used a Gaussian distribution in the GLM, while using for all other parameters a binomial distribution in the GLM. To reduce the skewness of the proportion of pair-flips during early hierarchy formation we applied a log(x + 1) transformation.

**Table 1.** Quantitative measures of dominance behaviour and network for the three versions of the model, namely the full model, the model without the spatial component, and without the winner-loser effect. The hierarchy is established using the average dominance index (ADI).

| *Measure* | **Full model** |  | **Without spatial component** |  | **Without winner-loser effect** |  | *P-value* |
| --- | --- | --- | --- | --- | --- | --- | --- |
|  | *Value* | *SE* | *Value* | *SE* | *Value* | *SE* |  |
| 1) Rank-changes (% of interactions) | 1.7%<sup>a</sup> | ±0.2 | 2.2%<sup>ab</sup> | ±0.2 | 2.4%<sup>b</sup> | ±0.2 | 0.01 |
| 2) Rank-changes correlated with pair-flips | 16.4%<sup>a</sup> | ±3.7 | 21.1%<sup>a</sup> | ±3.6 | 42.3%<sup>b</sup> | ±5.0 | < 0.001 |
| 3) Rank-changes during first 60 interactions | 54.6% | ±5.2 | 61.2% | ±4.5 | 58.8% | ±7.9 | 0.74 |
| 4) Hierarchy differentiation (steepness) | -0.29<sup>b</sup> | ±0.00 | -0.25<sup>a</sup> | ±0.00 | -0.30<sup>b</sup> | ±0.00 | < 0.001 |
| 5) Pair-flips (% of interactions) | 3.9%<sup>a</sup> | ±0.2 | 7.8%<sup>b</sup> | ±0.3 | 8.5%<sup>b</sup> | ±0.3 | < 0.001 |
| 6) Pair-flips during first 60 interactions | 9.1% | ±3.3 | 11.5% | ±1.6 | 14.2% | ±5.0 | 0.29 |
| 7) Bursts (% of interactions) | 52.3%<sup>c</sup> | ±0.6 | 28.3%<sup>a</sup> | ±0.5 | 48.3%<sup>b</sup> | ±0.6 | < 0.001 |
| 8) Maximum burst length | 11.2<sup>b</sup> | ±0.7 | 4.1<sup>a</sup> | ±0.1 | 10.3<sup>b</sup> | ±0.8 | < 0.001 |
| 9) Average linearity (proportion fully transitive states) | 0.98<sup>b</sup> | ±0.00 | 0.99<sup>c</sup> | ±0.00 | 0.96<sup>a</sup> | ±0.00 | < 0.001 |

The values in table 1 are averages and standard error (SE) measured over 14 runs. The last column shows the test results (GLM) of the effect of the model version on the respective response variable. Results from the post-hoc Tukey tests are shown as letters (a, b, c) in superscript. If a letter is not shared between two values this indicates a significant difference between these values. For instance, the proportion of rank-changes (item 1 in Table 1) is significantly lower in the full model than in the model without winner-loser effect. If a letter is shared, there is no significant difference between these groups, e.g. considering the proportion of rank changes, between the full model and that without the spatial component.

The minimal adequate model was determined by performing an analysis of variance (ANOVA) and continuing to drop the predictor variable with the highest p-value until all predictor variables left in the model had a p-value below the significance level of 0.05. In order to determine which model versions significantly differed from one another, we applied a Tukey test post hoc to the minimal adequate model of each measure.

Removing the spatial component or the winner-loser effect from the model had a significant effect on all measures shown in Table 1, except the proportion of rank-changes and pair-flips early in the formation (items 3 and 6 of Table 1) for which no minimal adequate model was found.

## Results

### 1. Behavioural dynamics

#### 1.1. Rank development

In hens the hierarchy was highly differentiated and stable. Rank changes were few and happened mostly during the first stage of hierarchy formation, for instance, a top-ranking individual often emerged early on and subsequently maintained its position. Rank changes in a pair were not preceded by a reversal in the direction of attack in that pair (a pair-flip, see next section for a definition), but were supposed to result from one of the members of the pair attacking others lower in rank with as consequence that it surpassed an individual ranking above itself.

Rank development in DomWorld (full model) is characterized by its stability (point 1 in Table 1) and a strong differentiation of the hierarchy (4 in Table 1). The top-ranking individual often (in 10 of the 14 runs, see appendix 2) emerged early in the run, maintaining its rank throughout the run. Rank changes that involved individuals at lower rank positions are distributed more evenly over time (Fig 4).

**Fig 4.**
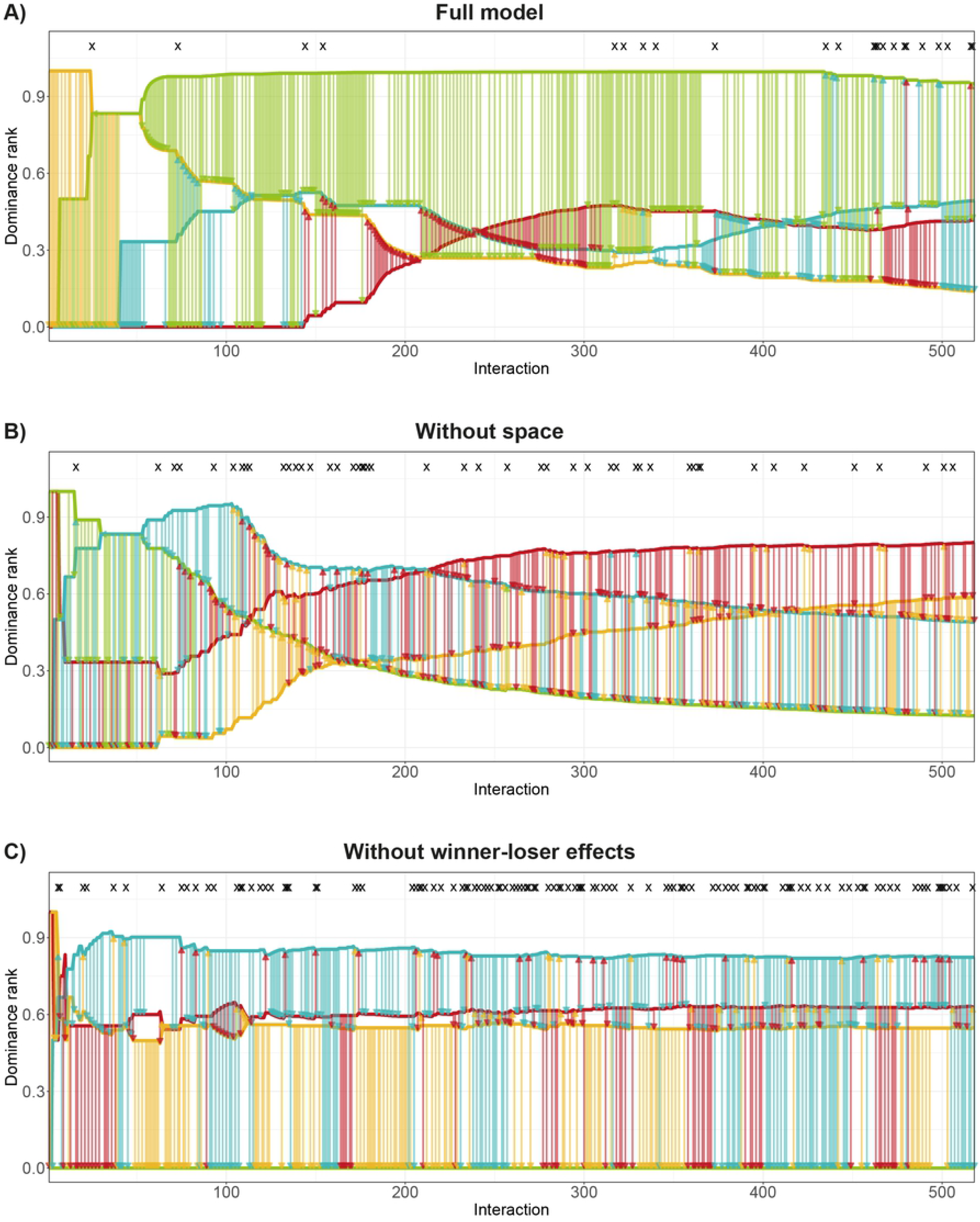
Music notation graph of rank development over interaction count for A) the full DomWorld model, and the DomWorld model B) without the spatial component and C) without the winner-loser effect. The horizontal lines represent the rank of each individual based on the average dominance index (ADI). The vertical arrows represent separate fights and point from the winner to the loser, in the colour of the winner. Pair-flips are marked with an ‘X’ at the top of the graph. Rank changes are shown as crossing horizontal lines.

Without a spatial representation or without the winner-loser effect the model leads to more changes of rank (1 in Table 1). Additionally, without a spatial representation when meeting others randomly, rank changes more often involve all rank positions over the entire length of the run (appendix 2), and result in a hierarchy that is less steep than in the full version of the model and the model without the winner-loser effect (4 in Table 1).

In all model versions more than half of the ascensions in rank occurred during the first 60 interactions (3 in Table 1). In the leadup to a rank change, the increase in the dominance value of the individual that will ascent in rank comes about by a combination of attacking lower ranking individuals and attacking the individual ranking immediately above itself with which it will swap rank, but not often by attacking others that are much higher in rank (Fig 4, also see appendix 2). Only one in six rank changes in the full model is directly preceded by a pair-flip (2 in Table 1). When removing the winner-loser effect from the model, rank changes more often correlate with a pair-flip than in the full model (2 in Table 1).

#### 1.2. Pair-flips

An interaction is classified as a pair-flip when a loser from an interaction wins from the same opponent in the subsequent fight. Hence a lower frequency of pair-flips indicates a more stable hierarchy. In hens, pair-flips were reported to be scarce, and half of them occurred during the first 60 interactions. Pair-flips were often quickly followed by another pair-flip indicating immediate retaliation of aggression.

In the full DomWorld model pair-flips occurred about half as often as in the model without space or without the winner-loser effect (5 in Table 1). In contrast to the pattern in hens, pair-flips in DomWorld (all versions) were not concentrated during early hierarchy formation (6 in Table 1) and were usually not directly followed by a counter pair-flip (Fig 4).

#### 1.3. Bursts

Attacks (pecks) by hens were often repetitive, involving one individual attacking the same opponent several times in a series. The duration of these ‘bursts’ followed a power-law with a maximum length of around 120 repetitive pecks. Pecks during bursts were directed down the hierarchy by all individuals except the lowest ranking individual. This is obvious because it does not have any individual lower in rank to attack.

About 50% of the interactions are part of bursts in the full model and in DomWorld without the winner-loser effect (7 in Table 1), with a maximum length of about 10-11 consecutive interactions (8 in Table 1). Without space in DomWorld, the percentage of interactions involved in bursts is halved and the maximum length of a burst reduces to approximately 4 interactions. In all model versions the average number of interactions in a burst was higher the greater the dominance of the attacker (Fig 5).

**Fig 5.**
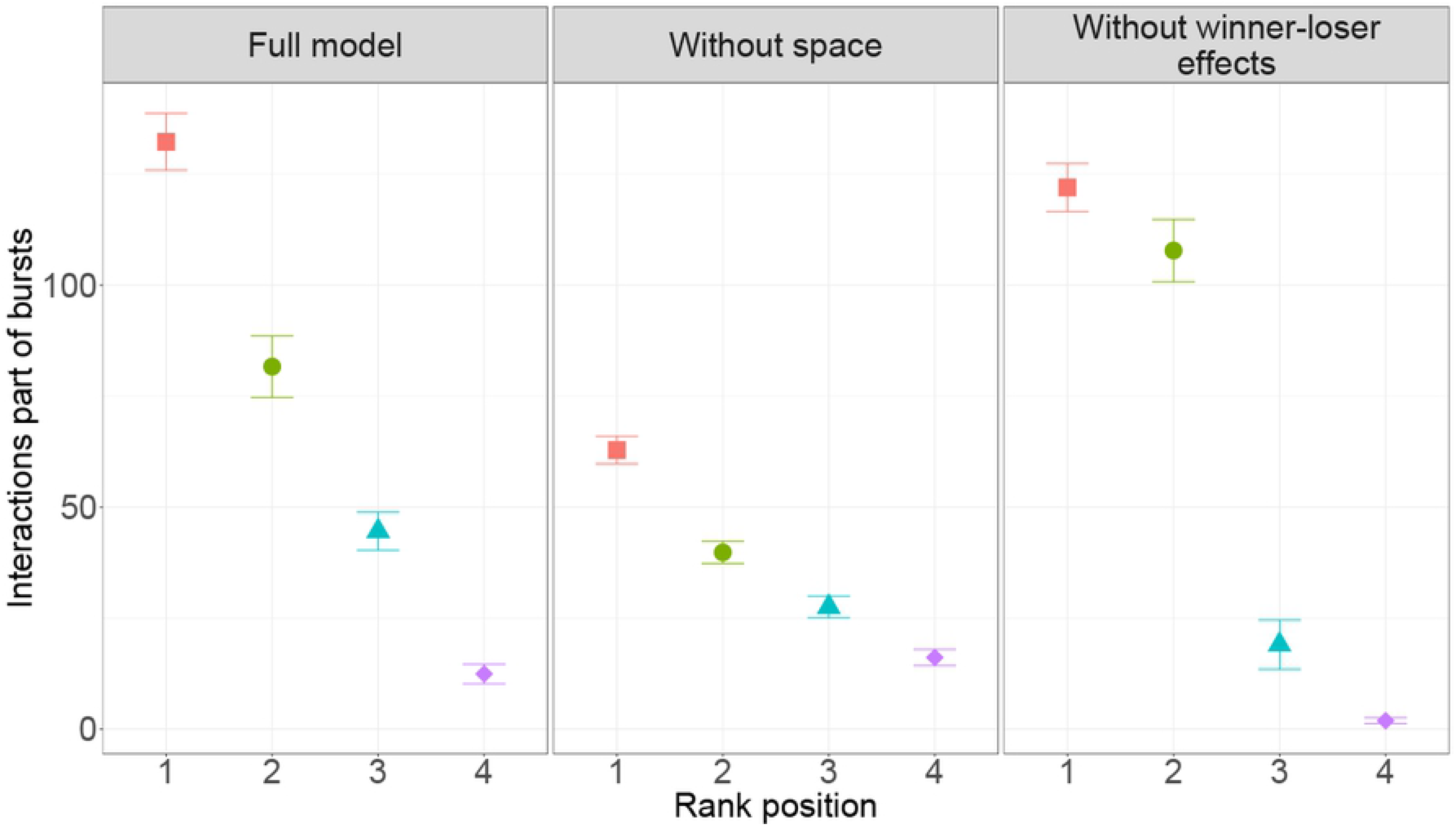
Average (and standard error, SE) number of interactions part of bursts per run. This is shown for each model version per rank position (rank 1 is most dominant).

#### 1.4. Spatial distribution

Even though a group size of 4 individuals is small, a spatial structure still emerges in which the dominant individual is more often in the centre of the group, while the lower ranking individuals are on average further away from it (Fig 6).

**Fig 6.**
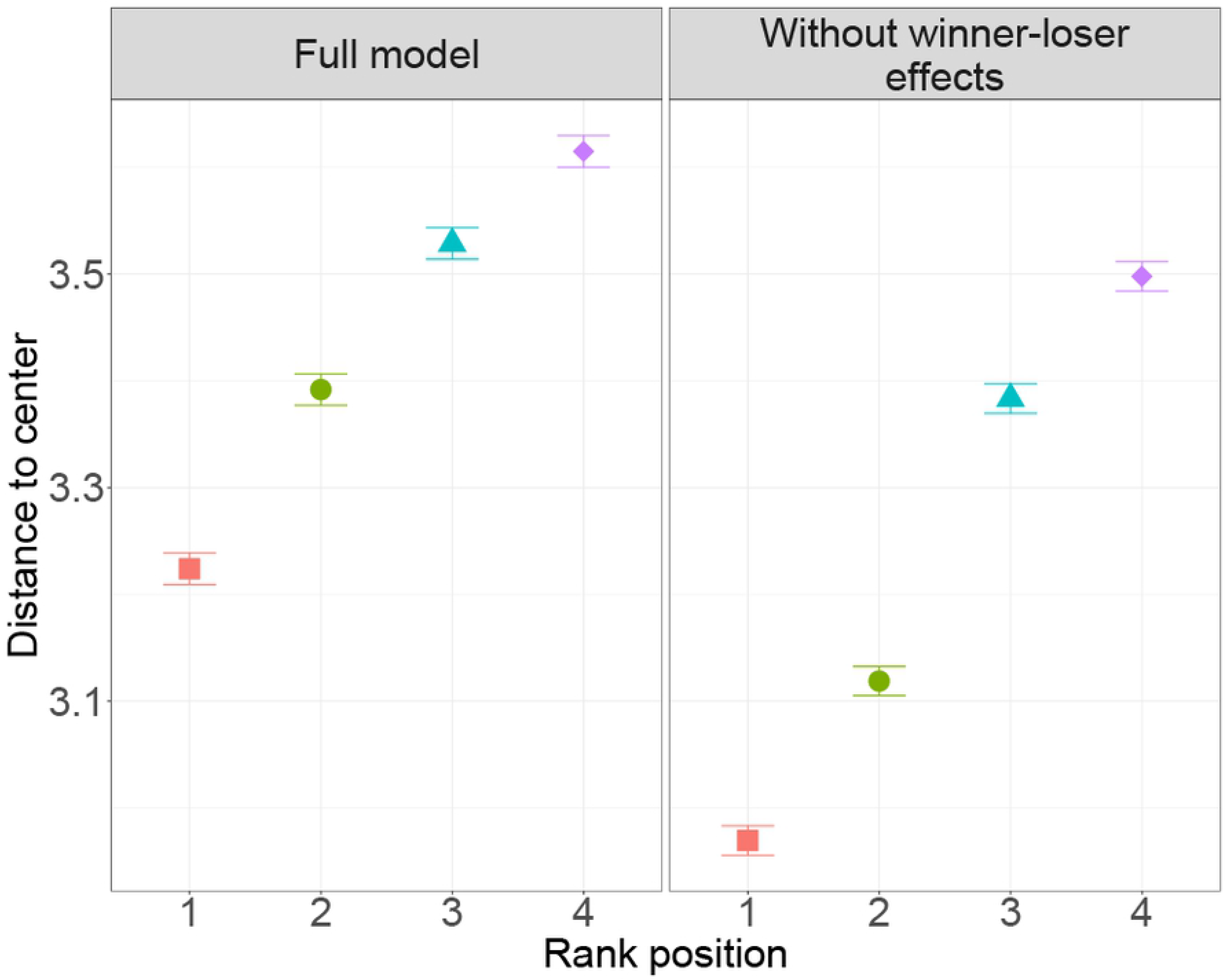
Average (and standard error, SE) distance to the centre of the group. This is shown per rank position (rank 1 is most dominant) for the full model and the model without the winner-loser effect.

### 2. Network analysis of triad motifs

To investigate the development of the dominance network as a whole, all its triadic combinations of individuals are assigned a triad motif. The collection of all triad motifs forms the state space of the network, see the section *Patterns of network structure* in *Methods*. The network of groups of hens developed rapidly via different paths until they reached a complete network with 6 links. Here those states occurred more often that either comprised of one or more transitive triads, no cyclic triads or one individual that is dominant over all others (DAO).

Similar to hens, the complete network of 6 links was reached fast in DomWorld. Less than one tenth of the total interactions resulted in a state with less than 6 links in each of the model versions (see N for each link-class in Fig 7). Also, the developmental path through the possible network states until a completely connected state (6-links) was reached, varied greatly among runs. Therefore, we will focus only on states with 6 links.

**Fig 7.**
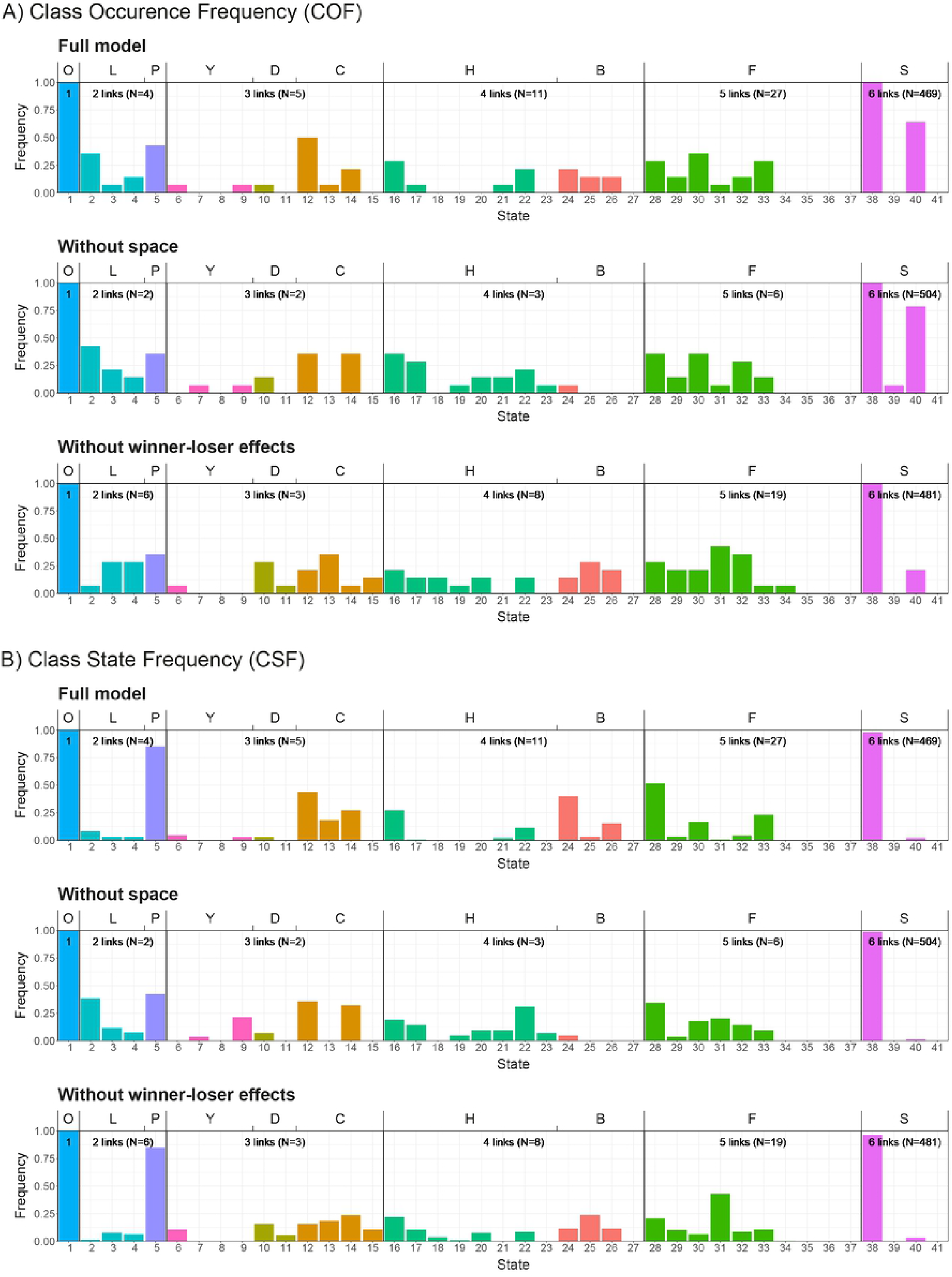
Histogram of A) in how many groups did a state occur at least once (Class occurrence frequency, COF) and B) how many times did a state occur over all runs relative to the total occurrence of states with the same number of links (Class state frequency, CSF). The state index (bottom), class letter (top) and colours correspond to those in Fig 3. Note that neither Class occurrence frequency, COF nor Class state frequency, CSF reflects the absolute number of the occurrence of a state. For each link-class the average number of interactions per run is shown as ‘N’ to give an indication of accuracy.

Of the 4 states with 6 links (Fig 3), state 38 is the only one that is fully transitive, with 4 transitive triads. State 39 and 40 comprise 1 cyclic triad and 3 transitive triads, and state 41 has 2 cyclic triads and 2 transitive triads. State 38 and 39 are the only states that have an individual that dominates all others (DAO), whereas state 38 and 40 contain an individual who is submissive to all others.

The frequencies of triad motifs in the original model and its two derived versions are similar (Fig 7B). Almost all (96-99%) interactions result in a network with state 38. The abundance of state 38 implies a high proportion transitive states, indicating the linearity of the hierarchy. The difference in frequencies of state 38 across the three model versions directly contributes to significant but small difference in their proportion of transitive states (9 in Table 1). Of the other states with 6 links, state 40 is the most common in all model versions (1-4%) and is together with state 38 the only state with 6-links that contains an individual that is submissive to all others. States 39 and 41 occur in is less than 1% of all interactions.

## Discussion

As to the similarity between aspects of the hierarchy in hens and in groups in the full model, DomWorld, we showed that in DomWorld, groups developed a highly linear and stable hierarchy that featured similar characteristics to those of hens (Table 2). After the value of the model parameter *intensity of aggression* was increased compared to former settings, that were relevant to macaques, the hierarchical linearity was similar to that in hens (11 in Table 2). Also the frequency of rank-changes, pair-flips and bursts in the model resembled those in hens (2, 5 & 7 in Table 2). The hierarchy in DomWorld developed rapidly whereby most changes in rank occurred early in the development of the hierarchy and soon most network states were fully connected with 6 links. The frequency distribution of the complete network states (with 6 links) in DomWorld resembled that of hens (Fig 8). A difference is that in DomWorld intransitivity is mostly the result of state 40, while in hens intransitivity comes from all states that contain one or more cyclic triads, thus also from states 39 and 41.

**Table 2.** Comparison between patterns of hierarchical development in the full model DomWorld and in groups of hens(16).

|  | **DomWorld**<br>*Full model* | **Hens**<br>*Lindquist and Chase (16)* |
| --- | --- | --- |
| 1) Hierarchy differentiation | Steep ($R = -0.29$) | Steep ($R \approx -0.29$) |
| 2) Rank changes per run | Few (8.7/run) | Few |
| 3) Rank position is determined rapidly | Yes (55% rank changes in first 60 interactions) | Yes |
| 4) Top-ranking individual often emerges early and consolidates its position while late rank changes occur in lower ranks | Yes (10/14 runs) | Yes |
| 5) Bursts (proportion of fights) | 0.52 | 0.49 |
| 6) Maximum duration of bursts | 17 interactions | ~120 interactions |
| 7) Pair-flips (proportion of fights) | 0.04 | 0.02 |
| 8) Percentage pair-flips during early development (< 60 interactions) | 9% | 51% |
| 9) Pair-flip is often quickly followed by a counter pair-flip | No | Yes |
| 10) Rank changes correlate with pair-flips | Few (16.4%) | No |
| 11) Average linearity (proportion transitive states) | 0.98 | ~0.98 |

**Fig 8.**
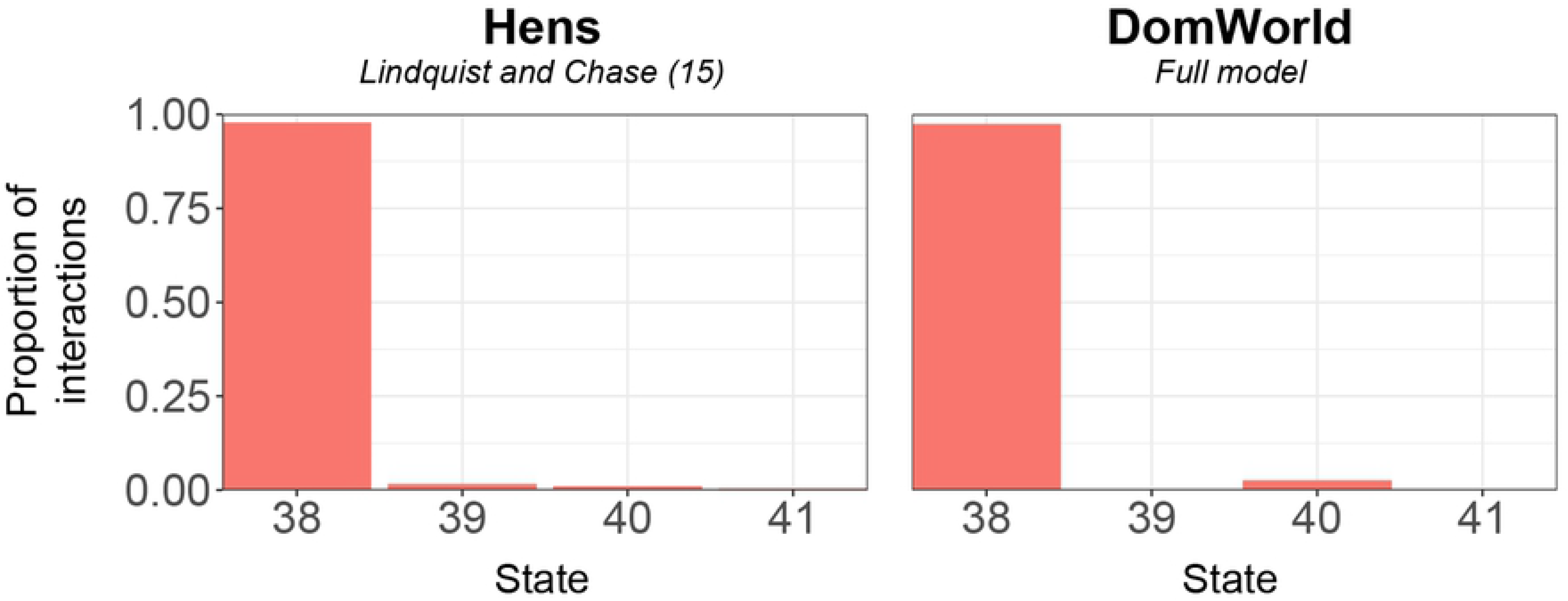
Comparison of the frequency of complete states (6 links) in the dominance network between the model DomWorld and hens as reported by Lindquist and Chase (16).

A few behavioural patterns in hens were absent in DomWorld. Most notably the long series of repeated attacks within a pair (6 in Table 2) and the immediate retaliation of aggression described for hens were absent (9 in Table 2). Further, pair-flips in DomWorld were more evenly distributed over the length of the run, whereas in hens they clustered during early hierarchy development (8 in Table 2). An explanation for these differences might lie in two methodological problems of matching the behaviour of individuals in our computational model to empirical data.

First, the observations of hens were different from those in DomWorld. In hens attacks predominantly consisted of pecks. In DomWorld dominance interactions represent complete fights, with an initiator that starts the fights, and a winner and a loser, where the loser flees after the fight. Possibly the lack of detail of fights in DomWorld prevents the occurrence of both long series of individual attacks (bursts) and the immediate retaliation of aggression by a lower ranking individual that were found in hens.

Second, the spatial environment of individuals in the model differs significantly from that of the groups of hens. The hens were confined in a cage (16, 27), whereas in DomWorld agents moved through an unlimited space (21). Thus in DomWorld the loser of a fight, often subordinate to the attacker, may flee without restriction, whereas in hens an individual is limited by the edge of the cage. Since this hinders the subordinate hen to flee its attacker, it suffers a longer series of attacks. The lack of escape options in hens may also result in a higher frequency of immediate counter aggression.

The lack of clustering in time of pair-flips in DomWorld as opposed to in hens may be due to the limits we set to dominance scores in the model. If dominance scores in DomWorld differ more between individuals this decreases the chance of a pair-flip. During hierarchy formation in DomWorld dominance scores were often restricted by these limits. Because all individuals started with the same dominance score which typically diverges over time, dominance score are less often limited during the initial part of the formation. Therefore, widening these limits in the model would mainly decrease the number of pair-flips later in the run and thus increase clustering of pair-flips during the first stage of hierarchy formation, as was reported for hens. Furthermore, asymmetrical clipping of the highest dominance scores might explain the relatively high frequency of state 40 in DomWorld compared to hens. This is because state 38 transforms to state 40 via a pair-flip that involves the highest ranking individual, whereas for state 38 to transform to state 39 it requires a pair-flip involving the individual with the lowest rank, and to transform to state 41 involves a pair-flip between both the lowest and highest rank (see Fig 3). Thus if the chosen limits more often restrict the dominance score of the lowest ranking individual than that of the highest ranking individual this may promote the number of pair-flips involving the lowest ranking individual.

Based on their review of the literature and a comparison among 3 well-known winner-loser models and observations of real hens, Lindquist and Chase (16) concluded that the winner-loser effect is not sufficient to explain development of linear hierarchies in groups of hens. The authors suggest that hens instead are aware of the hierarchy and their own position in it and are striving to keep the hierarchy linear. For this, the authors propose processes that are cognitively sophisticated, such as transitive inference and eavesdropping. These processes are absent in DomWorld.

Transitive inference, with which individuals fill in transitive relationships for unobserved relationships, has indeed been found in a wide variety of species, including cognitively simpler species such as hens and recently even insects (28). Where transitive inference was long thought to be the hallmark of human reasoning, the ability of simpler species to solve transitive-inference tasks begs the question whether the mechanism underlying transitive-inference-like behaviour is truly cognitively demanding (29). Yet while cognitively simpler explanations have been proposed based on reinforcement history (30, 31), experimental evidence is lacking (32–35).

However, it is unclear whether the task commonly used to measure transitive inference is directly relevant to the social context of real animals such as dominance relations in a group (29, 36). The vast majority of evidence for transitive inference in animals has been collected with the so-called N-term series task, wherein animals are first trained and then tested using transitive series of arbitrary stimuli such as colours, odours or shapes. A study that illustrates this question, by Takahashi and colleagues (37), finds that three species (tree shrews, rats and mice), which in other studies were shown to solve the N-term series task (38, 39), were not able to solve two inference tasks in social context, while a fourth species (capuchin monkeys) could.

In the present study we show that the winner-loser effect in combination with a socio-spatial component successfully reproduces many of the characteristics of hierarchy development in hens without the need for cognitively sophisticated processes, such as transitive inference. Thereby it forms a plausible alternative to assuming the need of transitive inference in dominance processes. Removing the winner-loser effect from DomWorld, thus representing fixed individual capacities of winning, reduces the resemblance of the model to interactions patterns in hens to 4 out of 12 patterns (11 patterns in Table 2 and one in Fig 8) compared to the full model. Furthermore, by experimenting with the presence of the spatial configuration of group members in the model we show that spatial interaction is essential for the formation of a highly linear and stable hierarchy.

On the other hand, even though the model DomWorld shows patterns of hierarchy development resembling those in real hens, it cannot prove the existence of similar processes in real animals. Although challenging, future research should determine to what extent the winner-loser effect shapes dominance hierarchies, a pioneering example is a study with novel statistical methods that provided the first evidence in a wild and uncontrolled population of primates (baboons) for the role of the winner-loser effect in the dynamics of the hierarchy (40).

In recent years a broader call has been echoed to investigate the development of social networks over time, arguing that for testing hypotheses relevant for selection, dynamics, development and evolution of social networks, it is necessary to include temporal dynamics and spatial constraints (41–44). Along these lines further research may focus on collecting time-series of data of development of the hierarchy in other species in order to determine whether the combination of the winner-loser effect and the socio-spatial structure can generally explain the formation of linear dominance hierarchies, also in species with different dominance styles than hens.

## Acknowledgements

We would like to thank Hanno Hildenbrandt for his work on the implementation of the model DomWorld in C++ that was used in collecting the data for this study.

## Supporting information captures

**S1 Appendix. Parameter settings DomWorld.**

**S2 Appendix. Full set of music notation graphs.**

